# Exhaustive exploration of the conformational landscape of small cyclic peptides using a robotics approach

**DOI:** 10.1101/338483

**Authors:** Maud Jusot, Dirk Stratmann, Marc Vaisset, Jacques Chomilier, Juan Cortés

## Abstract

Small cyclic peptides represent a promising class of therapeutic molecules with unique chemical properties. However, the poor knowledge of their structural characteristics makes their computational design and structure prediction a real challenge. In order to better describe their conformational space, we developed a method, named EGSCyP, for the exhaustive exploration of the energy landscape of small head-to-tail cyclic peptides. The method can be summarized by (i) a global exploration of the conformational space based on a mechanistic representation of the peptide and the use of robotics-based algorithms to deal with the closure constraint, (ii) an all-atom refinement of the obtained conformations. EGSCyP can handle D-form residues and N-methylations. Two strategies for the side-chains placement were implemented and compared. To validate our approach, we applied it to a set of three variants of cyclic RGDFV pentapeptides, including the drug candidate Cilengitide. A comparative analysis was made with respect to replica exchange molecular dynamics simulations in implicit solvent. It results that the EGSCyP method provides a very complete characterization of the conformational space of small cyclic pentapeptides.

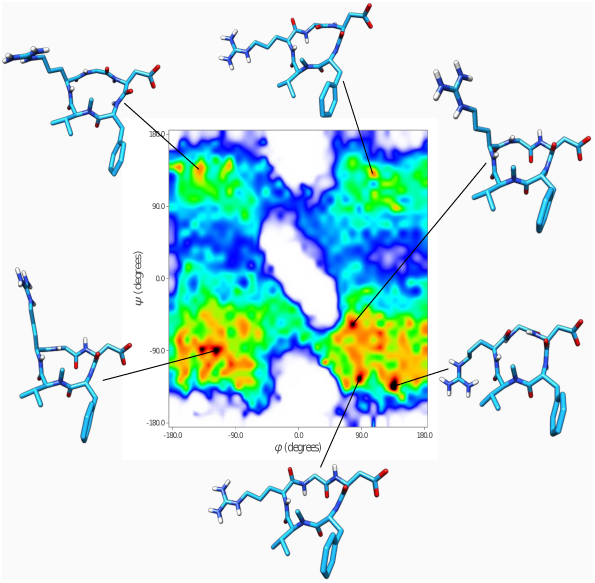

The paper presents a new method for the exhaustive exploration of the conformational energy landscape of small head-to-tail cyclic peptides. The approach is based on a multilevel representation of the peptide and the use of robotics-inspired algorithms.

## Introduction

After a relative decline for decades, peptides are turning back to the light as promising therapeutics drugs^1^. This is partially due to advances in chemistry to produce stable cyclic peptides as well as to the development of biotechnologies avoiding most of the drawbacks objected to peptides such as enzyme digestion or membrane penetration^2–6^. Peptides combine the advantages of proteins and small molecules: they have high selectivity as proteins, and metabolic stability, oral availability and low immunogenicity, as small molecules^2,6–8^. Compared to small ligands, peptides can occupy larger surfaces of interaction and reach higher specificities^2,6,7,9,10^. Thus, a particularly interesting application of peptides is the inhibition of protein-protein interactions involved in severe human diseases, including cancers and neurodegenerative disorders^11–13^. Therapeutic peptides are (re-)designed, aiming at improving their stability as well as their resistance to degradation by proteases. This can be achieved by cyclization and other chemical modifications, such as N-methylations or the use of D-amino acids^3,14–18^. Interestingly, combining cyclization and N-methylation has proven to improve membrane permeability, which is critical in the development of therapeutics against intracellular targets^7^. Finally, it has been shown that the N-methylation can modulate both the affinity and the specificity of a peptide for its target^19^. Therefore, cyclic peptides and their chemical modifications present pharmaceutical advantages that support the importance of their *in silico* design.

In spite of these promising properties in the field of pharmacology, we still are at the dawn of a sound understanding of cyclic peptides, in particular for head-to-tail cyclization, which would open the road to *in silico* predictions of their 3D structures. Indeed, there are only a few tens of structures annotated as “cyclic peptide” in the Protein Data Bank (PDB) and in the Cambridge Structural Database, as far as the length is less than 50 amino acids. Thus, template-based homology modeling methods are not adequate and only *ab initio* modeling is feasible. Besides, most of the *de novo* methods for modeling proteins are not adapted to cyclic peptides. Recent efforts have been made to develop or to adapt structure prediction tools to cyclic peptides. However, few of them are able to treat small head-to-tail cyclic peptides (with less than seven residues) involving chemical modifications. For example, PEP-FOLD^20,21^, PEPstrMOD^22^ and I-TASSER^23^ do not deal with sequences shorter than seven residues. The method “Simple Cyclic Peptide Prediction” of Rosetta^8^ can treat small peptides but cannot deal with N-methylation, so far. To our knowledge, only Peplook^24^ can deal with mixed-chiral small cyclic peptides and N-methylation. The difficulty to develop tools for predicting the structure of small cyclic peptides is essentially due to their very constrained structure that cannot contain any secondary structure nor hydrophobic core^15,25,26^, therefore with *ϕ* and *ψ* dihedral angles tending to fall outside the canonical allowed regions observed in the Ramachandran diagram^27^ for proteins^28^. In addition, the use of D-amino acids and N-methylations makes the modeling even more difficult^29^. To improve structure prediction, there is a real necessity of better understanding the conformational landscape of small cyclic peptides, and the related dihedral range needed to describe it. Our ambition was thus to explore the energy landscape of short, modified head-to-tail cyclic peptides. Indeed, while structure prediction aims at finding the most stable or probable conformations, global exploration methods are aimed to provide an overall picture of the conformational space.

Although molecular dynamics simulations and Monte Carlo methods can be used to explore the conformational space of linear peptides^30,31^, the application of these methods to cyclic peptides is less straightforward. This is mainly due to the ring-closure constraint, which leads to high energy barriers between the different meta-stable conformations^11,32,33^. Thus, it makes the conformational sampling very challenging. Actually, even for small systems, achieving a complete exploration of the energy landscape requires long simulation times when using basic approaches. Advanced methods are required to overcome this difficulty. For example, Replica Exchange Molecular Dynamics (REMD) simulations have been applied to cyclic peptides^34^. Metadynamics is a valuable alternative^11^, but the parametrization of the simulation, *i.e.,* the selection of the collective variables that are critical to capture the degrees of freedom (DoF) of the system, is still a bottleneck. Nevertheless, there is a need for alternative approaches more adapted to cyclic peptides than REMD or metadynamics for an unambiguous complete exploration of the conformational landscape, getting rid of the difficult assessment of convergence necessary in the various methods based on molecular dynamics.

Numerous methods have been proposed since the the seminal work of Go and Scheraga^35^ to efficiently sample cyclic molecules. We can mention for instance the work of Wu and Deem^36^, and of Coutsias *et al*.^37^. Inspired from these methods, we present a robotics-based approach for exploring the conformational landscape of head-to-tail cyclic peptides possibly involving mixed-chirality and N-methylated residues. Our method, called Exhaustive Grid Search for Cyclic Peptides (EGSCyP), is based on a multi-level representation of the peptide, and on the application of different algorithms at the various levels. Backbone conformations are first exhaustively sampled considering dihedral angles as the main variables. An inverse kinematics (IK) algorithm^38^ is used at this level to enforce loop closure. Then, for each backbone conformation, side-chains are placed and local minimization is performed using an all-atom representation. In this paper, we focused on pentapeptides, but our method also applies to tetrapeptides and can be trivially adapted to hexapeptides provided a simple parallelization of the algorithm.

In order to validate this approach, we sampled the conformational landscape of a set of three cyclic pentapeptides described in the literature^39,40^, containing the widely studied RGD motif. One of these peptides is Cilengitide^41^, which is an example of promising N-methylated RGD cyclic pentapeptide, developed by the Kessler group as inhibitor of protein-protein interaction of the *α*V*β*3 and the *α*V*β*5) integrins. It has reached the phase III of clinical trial for glioblastomas and is also under evaluation for other types of cancer^42^. This pentapeptide was chosen as a proof of concept since one structure is deposited and it is well studied experimentally. To evaluate the completeness and the accuracy of the EGSCyP approach, the results on these three cyclic peptides were compared with those obtained with REMD simulations.

## Methods

### Overview

EGSCyP applies a kind of “divide and conquer” paradigm. The global conformational exploration problem is divided into several sub-problems, each of which involves different variables. The backbone dihedral angles are treated first, since they are the most important degrees of freedom of a peptide. Among the backbone dihedral angles, the *ω* angles (corresponding to peptide bonds) are particularly rigid. Therefore, and aiming to reduce the combinatorial complexity, their values are randomly sampled from a Gaussian distribution centered at 180°, rather than systematically explored. The exhaustive exploration focuses on the *ϕ* and *ψ* angles. The loop-closure constraint imposes a non-linear relationship between these angles. More precisely, the value of 6 angles is determined from the value of the other *n* – 6 angles (*n* representing the total number of *ϕ* and *ψ* dihedral angles), using an inverse kinematics (IK) solver. In our approach, we assign these 6 dependent variables to the *ϕ* and *ψ* angles of three consecutive residues. Therefore, for a pentapeptide backbone, the remaining (independent) variables to be sampled are the *ϕ* and *ψ* of only two residues. Thanks to the low dimension, these four variables can be sampled with high resolution using an exhaustive grid search. The conformation of the side-chains is then sampled for each backbone conformation satisfying loop closure and without significant steric clashes. We have investigated two different approaches for solving this second sub-problem. Finally, the whole conformation is locally minimized at an all-atom level (i.e. considering all the degrees of freedom simultaneously). The multi-level model and the different stages of the EGSCyP algorithm are explained with more detail below.

### Molecular model

The representation of molecules is generally based either on the Cartesian coordinates of all their atoms, or on the set of internal coordinates corresponding to the relative positions of their covalently bonded atoms. This second representation can be defined by three types of DoF: bond lengths, bond angles and dihedral angles. It has been shown that the first two present low variations at room temperature^43^. Therefore, the model can be simplified considering these parameters as constants, adopting the rigid geometry assumption^44^, which means that the only DoF are the dihedral angles. This representation of a peptide allows modeling it as an articulated mechanism, similar to a robotic arm^38^ (see Figure 1). Thanks to this choice of representation, algorithms from robotics can be applied to explore the conformational space of molecules^45–47^. The approach we present in this paper uses both models: (1) the mechanistic one for the global exploration of the backbone conformation and the side-chain placement, (2) the Cartesian all-atom model for the refinement with the relaxation method.

**Figure 1:**
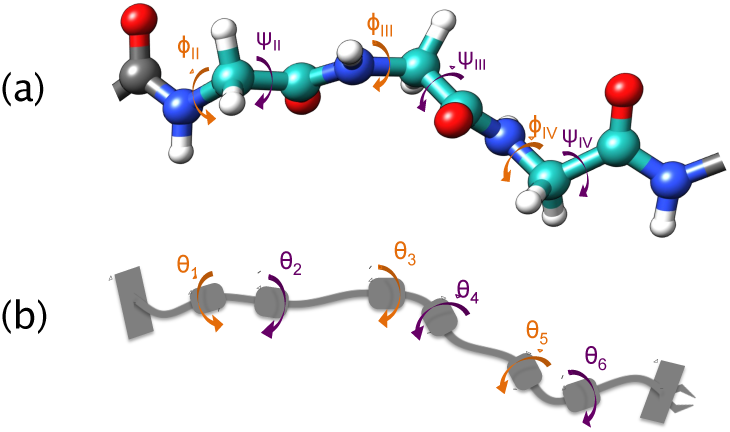
Analogy between a tripeptide (a) and a robotic arm (b). The dihedral angles *ϕ* (orange arrows) and *ψ* (purple arrows) of the tripeptide (in cyan) correspond to the revolute joints of the kinematic chain in this analogy.

### Sampling algorithm

The EGSCyP algorithm is summarized in Figure 2 for a pentapeptide. The exploration starts by the selection of two consecutive residues to be sampled, involving two pairs of *ϕ* - *ψ* angles. By default, the selected residues are the first and the last ones in the PDB file given as input (see dataset paragraph). In the following, these two residues are named *I* and *V*, respectively. The effect of the choice of the two sampled residues has been evaluated by testing all the possible positions (see results section). The sampling of all possible combinations of the two pairs of *ϕ* - *ψ* dihedral angles is made with a constant step size of 10° over the range from –180° to 180°. Then, for every combination, the five *ω* angles are randomly selected from a Gaussian distribution with a standard deviation of 10° centered around 180°. Then, the IK method is applied to close the cycle between the sampled residues (see below for more details). In other words, the IK method finds feasible values for the *ϕ* - *ψ* angles of residues *II*, *III* and *IV.* If there is no solution, the *ω* angles are sampled again until a solution is found or until a maximum number of iterations is reached (100 iterations in our implementation). If no solution is finally found, the next combination of torsion angles is tested. Otherwise, for each solution of the IK method, possible collisions between the backbone atoms (and the methyl carbon of N-methylated residues, if there was one) are checked. A collision is detected if the distance between two non-bonded atoms is less than 60% of the sum of their van der Waals radii^48^, thus accepting a small overlap. This geometric filtering was not chosen too restrictive, because the final relaxation could avoid small collisions not filtered at this step. Then, in absence of collision, the side chains are positioned using one of the following methods: (1) the SCWRL4 software^49^, or (2) a variant of the Basin Hopping (BH) method^50^. Details about these methods are described below in the paragraph about side chains placement. Finally, a few steps of structural relaxation are performed using Sander from AmberTools 16^51^. This last stage allows releasing constraints imposed by the rigid geometry representation. The relaxation is made with the amberff96 force field^52^ with a Generalized Born implicit solvent (all default parameters were set on). The choice of this force field and implicit solvent was made based on the reasonable performance of this combination on peptides compared to other force fields^53^. During relaxation, the *ϕ* and *ψ* dihedral angles from the two sampled residues are restrained to their sampled values. For the calculations presented below, the maximum number of cycles of minimization was set to 1000, with 500 steps of steepest descent followed by conjugate gradient. The convergence energy criterion for the minimization was set to 0.1 kcal/mol-Å. At the end of each iteration, for each solution, a conformation is built (*i.e*., atomic coordinates are extracted) and the dihedral angles values are recorded for all the residues as well as the energy of the peptide, calculated by Amber. A connectivity graph is created during the sampling process, with one node for each conformation. In the sampling grid, neighbor nodes are linked in such a way that each node is connected to all the nodes having the four sampled dihedral angles within a window of ±10°. As explained below, the connectivity graph is used within the side-chain placement method.

**Figure 2:**
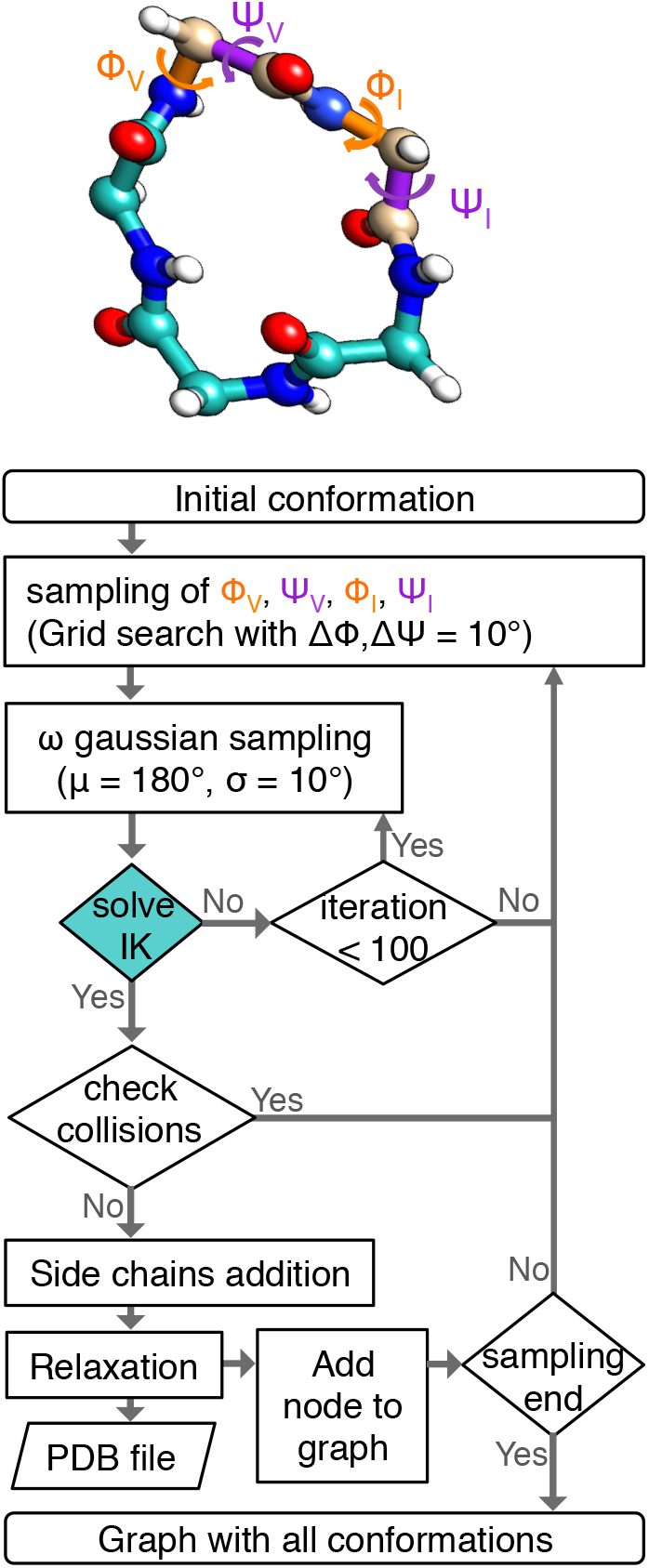
Flowchart of the EGSCyP algorithm. The backbone of a pentapeptide is represented to illustrate the sampled dihedral angles *ϕ* and *ψ* colored in orange and purple respectively. The tripeptide on which the IK is applied is colored in cyan.

### Inverse Kinematics

As previously described, a molecule can be modeled as an articulated mechanism. Using standard conventions applied in robotics, such as the *modified Denavit-Hartenberg* convention ^54^ used in this work, a Cartesian coordinate system *F*_*i*_ is attached to each rigid body, which corresponds to a small group of bonded atoms. Then, the relative location of two consecutive frames in the chain can be defined by a homogeneous transformation matrix ^*i*–1^*T*_*i*_. Assuming constant bond lengths and bond angles, the only variable parameter in this matrix corresponds to a bond torsions, *θ*_*i*_. In the case of a peptide backbone, and if we assume that the *ω* angles are also fixed to a given (sampled) value, the variables are the *ϕ* and *ψ* angles. Thus, as illustrated in Figure 1, the backbone conformation for a tripeptide composed of residues *II*, *III*, *IV* is defined by the vector of six angles {*θ*_1_*, θ*_2_*, θ*_3_*, θ*_4_*, θ*_5_*, θ*_6_} = {*ϕ*_*II*_, *ψ*_*II*_, *ϕ*_*III*_, *ψ*_*III*_, *ϕ*_*IV*_, *ψ*_*IV*_}. The IK problem relies in finding values for this vector of angles such that they solve the following matrix equation: ^0^*T*_1_^1^*T*_2_^2^*T*_3_^3^*T*_4_^4^*T*_5_^5^*T*_6_ =^0^ *T*_6_, where ^0^*T*_6_ is the homogeneous transformation matrix representing the relative pose of the first frame *F*_0_ and the last frame *F*_6_. In our case, *F*_0_ and *F*_6_ are determined from the conformation of the other two residues in the cyclic peptide, which were exhaustively sampled as explained in the previous section.

For solving the IK problem, we apply the method proposed by Renaud^55,56^. This semi-analytical solver, based on algebraic elimination theory, is rooted in the work of Lee and Liang^57,58^. Our implementation incorporates ideas proposed by Manocha and Canny^59^ to improve numerical robustness. The solver is very computationally efficient, requiring about 0.1 milliseconds on a single processor. It was successfully applied in previous works on protein and polymer modeling^60–62^. Note however that our approach does not rely on a particular IK solver. Other methods available in the literature could be applied (e.g.^37,59,63^). Nevertheless, we recommend the application of (semi-)analytical methods, which, in the present application case, have several important advantages with respect to purely numerical approaches such as CCD^64^. The main advantage is that they simultaneously provide all the solutions to the IK problem: up to 16 for an articulated mechanism with six revolute joints, as our tripeptide model. In addition, they provide the exact solution in a single iteration, not suffering from slow convergence issues.

### Side-chain placement

Two methods were implemented and evaluated for the side-chain positioning. The first one was SCWRL4^49^, a widely-used and efficient tool for the prediction of side-chain conformations, based on a rotamer library. In our case, the applicability of this method is limited, because it cannot deal with the N-methylated residues (it replaces them by glycines). D-amino acids are not represented in the rotamer library either. Moreover, SCWRL4 does not deal with cyclization: collisions usually happen between the side-chains of the first and the last residues. Within our multi-stage exploration approach, the subsequent all-atom relaxation (performed with Amber) can solve some of these problems. However, wrong side-chain conformations may remain. Finally, despite the fairly high speed of SCWRL4, its use as a third-party program requiring system calls slows down the overall computing time.

The second approach used for the side-chain placement is much more general. It is based on stochastic optimization methods. More precisely, we applied a variant of the Monte-Carlo-minimization method^65^, also known as Basin Hopping (BH)^50^, as explained in previous work^66^. This global optimization method consists of iteratively applying a random structural perturbation (global search) followed by a local energy minimization. At each iteration, the new local minimum is compared with the one generated in the previous iteration, and it is accepted or rejected based on the classical Metropolis criterion. In our implementation, the structural perturbation was applied to a randomly selected number of *χ* angles in a first step. Then, the local minimization based on a simple Monte-Carlo (MC) method at very low temperature (in order to remain in the local basin), applied small random perturbations to all the *χ* angles. Although this type of local minimization is in general less computationally efficient than gradient-based approaches usually applied within BH, it is also less sensitive to small local traps. It requires about 10 seconds on a single processor (it is the most expansive step of our algorithm).

In order to decrease the overall computational cost, the BH method was not systematically applied to place the side-chains for every sampled conformation of the peptide backbone. When a neighboring conformation of the peptide (*i.e.*, a neighbor in the connectivity graph) has already been generated by the exhaustive exploration algorithm, the side-chains conformation was then generated by local minimization from this neighbor, using the local Monte-Carlo search. The computational time is reduced to 10 milliseconds and is equivalent to the use of SCWRL4. Energy evaluation within the side-chain placement procedure was performed using the same force field as in the subsequent relaxation process: the amberff96 force field with a Generalized Born implicit solvent.

### REMD

For comparison, Replica Exchange Molecular Dynamics simulations were performed with GROMACS 5.1.2^67^. As in the EGSCyP method, the simulations were performed using the OBC (Onufriev, Bashford, and Case)^68^ GBSA implicit solvent model and the Amber96 force field^52^. A short minimization was first done with an alternation of one step of steepest descent step every 500 conjugate gradient steps. The maximum number of steps was set to 50,000 with a step size of 0.01 nm and an energy convergence criterion of 10 kJ/mol.nm. The thermalizations and simulations were made with eight replica varying in temperature from 300K to 450K (at 300K, 313K, 329K, 347K, 367K, 391K, 418K, 450K). The temperatures of the replica were chosen to keep constant the probability of accepting exchanges. Only a short phase of thermalization was realized on 100 ps (50 000 steps). Then, the REMD simulations were performed with parameters inspired from the work of Wakefield *et al*.^34^. The exchanges between neighboring replicas were made every 10 ps. Each replica ran for 2.4μs, yielding a total simulation time of 19.2 μs per peptide. The atomic coordinates were recorded every 10 ps. However, contrary to the work of Wakefield *et al*.^34^, we did not change the torsion scaling parameters to lower the *ω* angle torsional barriers and to accelerate the cis/trans sampling of N-methylated residues. We kept it to the default value, because the cis conformations were only populated at a few percent in the study of Wakefield *et al*.^34^.

### Clustering

In order to rapidly and easily identify the main energy minima produced by EGSCyP and REMD simulations, we developed a simple clustering method inspired from the one recently proposed by Hosseinzadeh *et al*.^25^. It is based on the energy of conformations and the Root Mean Square Deviation (RMSD) calculated upon the *ϕ*-*ψ* dihedral angles^69^. For EGSCyP, the energy taken into account is the final energy computed by Amber at the end of the relaxation. For REMD simulations, the potential energy for each frame of the lower-temperature replica is computed with Gromacs. The global energy minimum is selected as the center of the first cluster. Then, the RMSD between this minimum and all the other conformations is computed. All the conformations presenting a RMSD inferior to a threshold are put into the first cluster. The choice of this threshold will be discussed below, together with the results. Next, the same procedure is applied again: among the remaining conformations, the structure with the lowest energy is selected as the center of the second cluster and the RMSD is again computed for creating the second cluster. The procedure is repeated until all the conformations were included into a cluster. This approach, unlike classical clustering based only on distances, may be biased toward low-energy basins. However, its application in the context of this study to find and compare the energy minima basins seems quite appropriate. As a final stage, the representative structures of clusters from both methods are minimized without any constraint with Amber in order to be comparable.

### Dataset

In this study, we considered three RGD cyclic pentapeptides selected from the work of the Kessler group^39,40^. The sequences of these peptides are presented in Table 1. For both the REMD simulations and EGSCyP method, all the cyclic structures were generated using UCSF-Chimera^70^. This tool allows modeling non-natural amino acids, such as D-amino acids or N-methylated ones. The cyclization was made head-to-tail, *i.e.*, a peptidic bond was created between the N-ter and the C-ter amino acids. The exhaustive exploration process performed by EGSCyP is independent of the initial conformation. The topology files were created with Tleap from AmberTools 16^51^, using the Amber ff96 force field^52^. The Amber topologies and partial charges of the N-methylated residues were computed with the RED-Server^71^. For the REMD simulations, the Amber topology files were converted into Gromacs topology files with Acpype^72^ and manually modified for compatibility with recent versions of Gromacs.

**Table 1:** Dataset of cyclic pentapeptides. Lower case letters indicate D-amino acids and single quote are for the N-methylated ones. RGDfV’ corresponds to Cilengitide (PDB ID: 1L5G).

| Peptide sequence | Type |
| --- | --- |
| RGDFV | natural |
| RGDfV | D residue |
| RGDfV' | D residue + N-methylated residue |

## Results and Discussion

### Exhaustive grid search

We used the EGSCyP method for the three peptides presented in Table 1. The level of exploration of the conformational landscape resulting from the EGSCyP approach was compared to the one produced by REMD simulations, over 2.4 μs for each replica of each peptide. Due to the low number of DoF of a cyclic pentapeptide and the relatively long simulation time, the REMD sampling is sufficiently exhaustive to be compared with the one obtained with EGSCyP (the convergence of the simulations and the percentage of accepting exchange between replica were verified; see supplementary material Figures S1, S2 and S3).

#### Choice of the sampled residues

We did a systematic test to verify that the choice of the two exhaustively sampled residues does not significantly affect the results of the EGSCyP method. It consisted of repeatedly applying the EGSCyP method, selecting each time a different combination of two consecutive residues to be exhaustively sampled. The test was applied to the RGDfV peptide (see Table 1 for nomenclature). The obtained landscapes are fairly similar, independently from the residues chosen to be sampled (detailed results are available in supplementary materials in Figure S4 and Table S4). The differences can be explained by four reasons: (1) the more exhaustive sampling performed for two pairs of *ϕ* and *ψ* angles (with respect to the angles solved by IK), which enforces the exploration of high-energy areas in the corresponding projections; (2) the randomized sampling process of the *ω* and *χ* angles; (3) the resolution of the discretization (*i.e*., 10° step size in the grid search) for the exploration of the two pairs of *ϕ* and *ψ* angles; (4) the imposed restrain on these four angles during the relaxation. The first of these reasons explains why the percentage of coverage of the Ramachandran diagrams (defined as the percentage of *ϕ*,*ψ* pairs, within the grid spacing of 10°, for which at least one conformation was found by EGSCyP) varies from 85% to 95% for the residues solved by IK, while it grows from 97% to 98% for the two exhaustively sampled residues. In spite of these variations, the low energy basins still remain in the same areas of the Ramachandran plot. The main differences are localized in areas of high energy. Therefore, we can make the simplification of sampling any two successive residues and still obtain a very complete landscape.

Table 2 shows a summary of the results obtained with EGSCyP that are detailed on the next paragraphs.

**Table 2:**
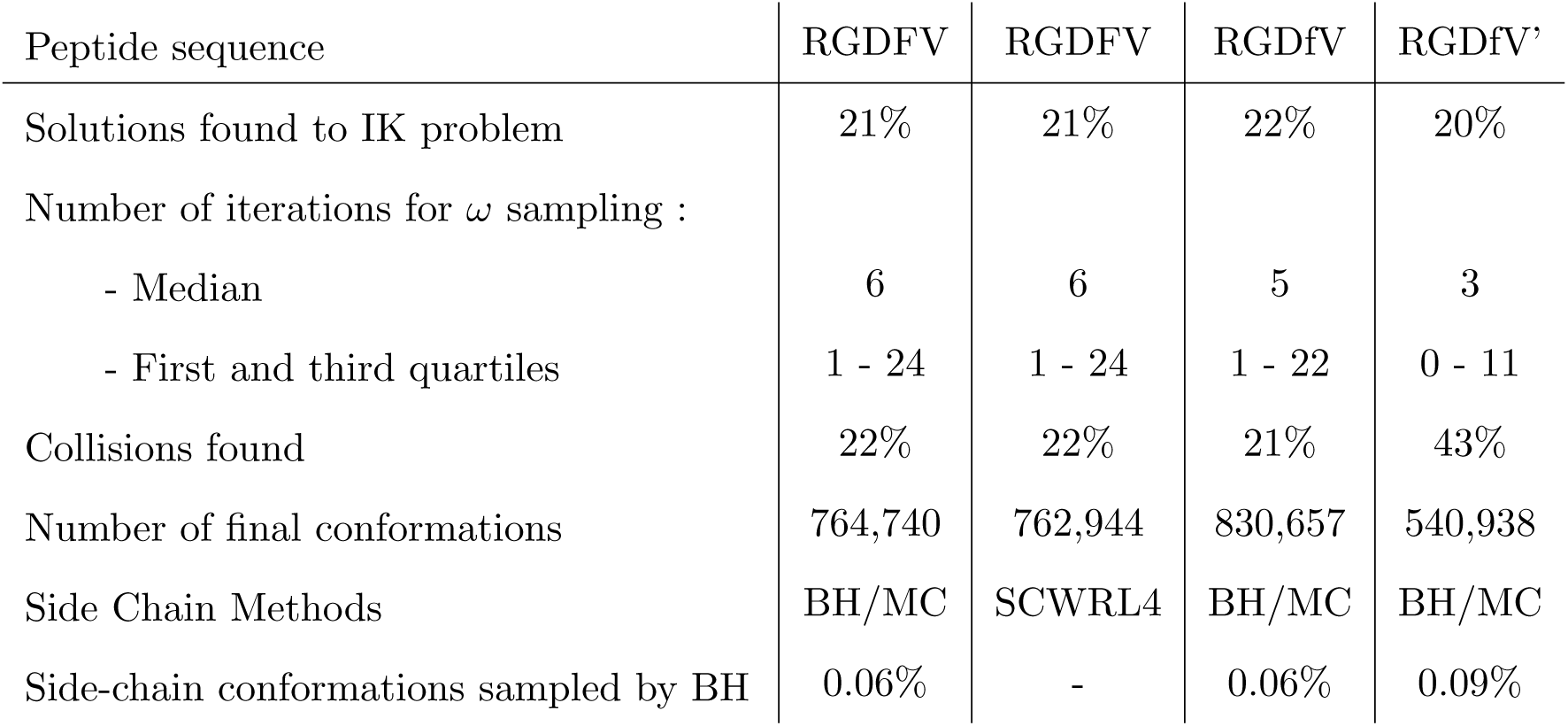
Summary of the performances of the EGSCyP method on the three cyclic pen-tapeptides. The RGDFV peptide is represented twice because of the two different methods used for side chain placement: SCWRL4 and the alternation of Basin Hopping (BH) and local minimization by Monte Carlo search (MC). The number of solutions of the IK problem is represented as a percentage of the cases with at least one solution among all the combinations of *ϕ*/*ψ* angles. The number of iterations for the *ω* angles sampling correspond to the number of iterations before the IK solver found a solution (until 100 iterations, in which case the *ϕ*/*ψ* combination is rejected). The percentage of collisions corresponds to the percentage of conformations among all the IK solutions containing overlapping atoms.

#### IK Solutions

The sampling of the four *ϕ* and *ψ* dihedral angles was realized with a step size of 10°, which means that (360/10)^4^ = 1.68 · 10^6^ combinations were tested for each peptide. For each *ϕ*, *ψ* combination, up to 100 combinations of the five *ω* dihedral angles were tested. 20%-22% of the 1.68 · 10^6^ *ϕ*, *ψ* combinations (*i.e*, about 350,000 combinations) yielded at least one solution to the IK problem. This proportion is rather constant among the different peptides, which is due to the fact that, at this step, the side chains are not taken into account and the number of accepted conformations is thus independent from the sequence. Indeed, the results among these various sequences differ only by (1) the geometry of the initial PDB structure, with the bond length and angles that can slightly vary between the initial structures (remember that they are kept fixed during the *ϕ* - *ψ* - *ω* sampling and the use of IK), and (2) a random factor which can be fixed by the use of a unique random-seed for the sampling of the *ω* dihedral angles. The fact that less than 1/4 of the *ϕ*, *ψ* combinations yielded IK solutions emphasizes the difficulty of ring closure for small cyclic peptides.

#### ω sampling

The median and the third quartile of the number of iterations over the combinations of the five *ω* angles sampling processed before a submission to the IK solver vary from 3 to 6 and from 11 to 24 respectively. Because solutions were generally found rapidly after a few iterations, the limit of 100 iterations is sufficient to find a solution, if any.

#### Number of collisions

The number of collisions between backbone atoms is rather constant (around 20% of the solutions found by IK per peptide) except for Cilengitide (RGDfV’). For this last peptide, the number of collisions increased up to 43%. This is the consequence of the N-methylated valine residue which very easily enters in collision with the backbone due to the small size of the cyclic pentapeptide. This illustrates the fact that N-methylations constraints the structure of peptides and reduces their conformational landscape.

#### Side-Chain placement

Among the three considered pentapeptides, RGDFV is the only one involving no chemical modification. Therefore, we used this molecule to compare the performance of SCWRL4 and the BH method (see below). The combination of BH and local minimization from a neighbor conformation was applied for the two other peptides of Table 1 with side chains. Note that the percentage of conformations for which BH was applied was very low. For a large majority of the conformations (around 99%), the side chains were placed by local minimization from a neighbor conformation in the connectivity graph. This shows that the graph is very dense, thanks to the exhaustiveness of the exploration.

#### Number of final conformations

The total number of conformations generated for each peptide is around 800,000, at the exception of Cilengitide (RGDfV’) for which only 540,938 conformations were found (because of the collisions mentioned above). Note that the number of conformations is significantly larger than the number of sampled combinations of *ϕ* - *ψ* angles (around 350,000). The reason is that, in most of the cases, the IK solver finds several conformations for the tripeptide. Indeed, two IK solutions were found for around 50% of the combinations for which the closure constraint can be satisfied. Excepting one case, the IK solver found at most up to eight conformations for the tripeptide. The exact numbers of solutions found for each peptide are presented in supplementary materials (Table S3).

### Validation on Cilengitide

#### Comparison of Ramachandran diagrams

The energy landscapes obtained with EGSCyP and REMD simulations were first compared using Ramachandran diagrams for each peptide and method. In Figure 3, the first two columns correspond to the conformations obtained by EGSCyP for the peptides RGDFV and RGDfV. The last two columns correspond to the maps for Cilengitide (RGDfV’) with the results obtained by the two methods: EGSCyP on the third column and REMD on the last column. Each line corresponds to one of the five residues of the cyclic pentapeptide. The diagrams corresponding to the comparision between EGSCyP and REMD for RGDFV and RGDfV peptides are available as supplementary material in Figure S5.

**Figure 3:**
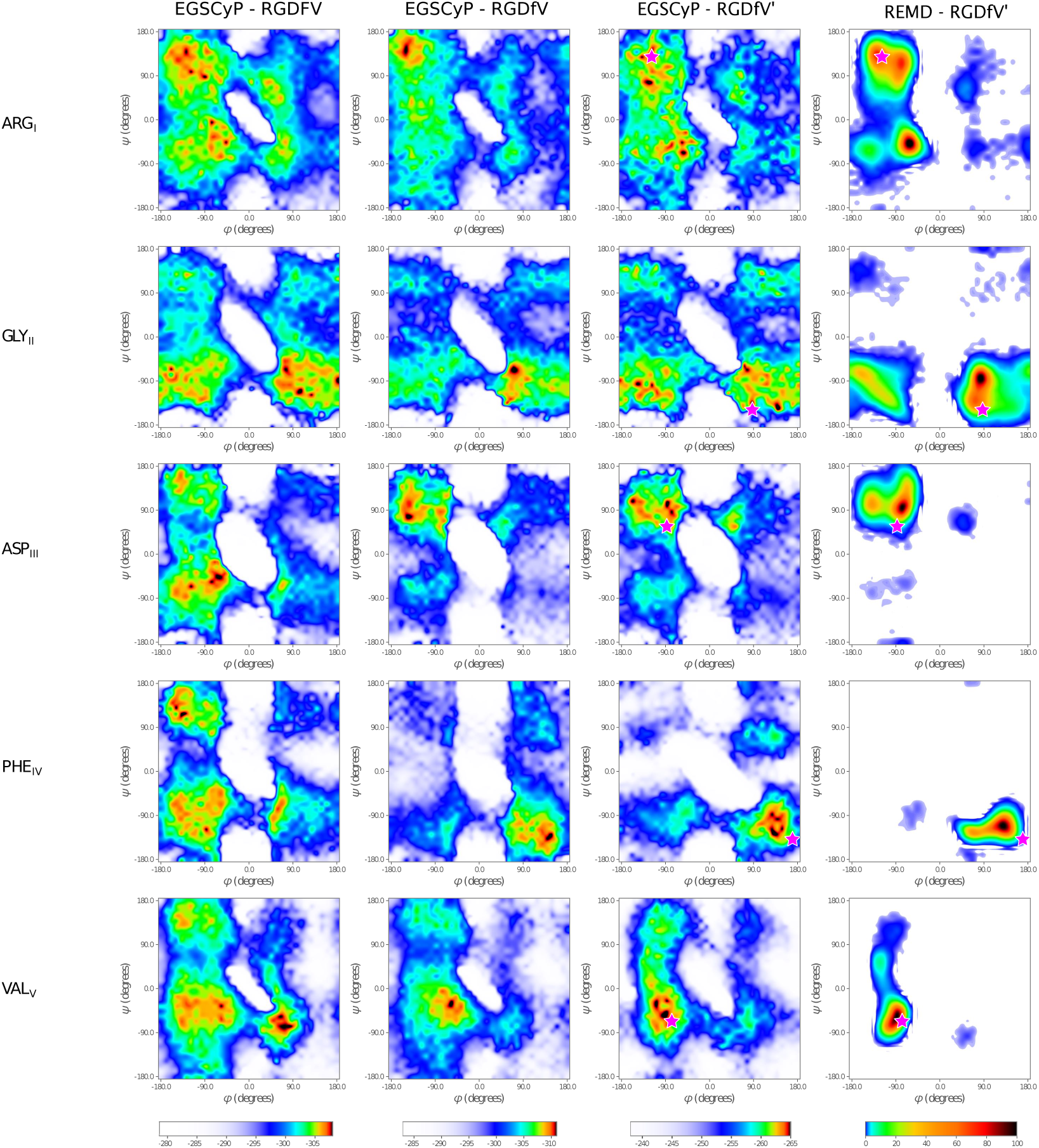
Ramachandran plots presenting the results obtained with the EGSCyP method for the RGDFV, RGDfV and RGDfV’ peptides (three left columns) and the REMD simulations for RGDfV’ (right column). The diagrams were created using the matplotlib python library^78^. For EGSCyP, the energy landscape was projected as a function of the dihedral angles *ϕ* and *ψ* for each residue (on each line). The color code corresponds to the minimal potential energy in kcal/mol of the whole peptide associated to each combination of *ϕ* - *ψ* angles. Energy minima are represented in black/red, while areas with no conformation are in white, and areas of high energy are in blue. For REMD, the normalized frequency 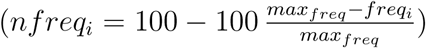 of the conformations found during the simulations at the lowest temperature replica is projected as a function of *ϕ* and *ψ*. The color code corresponds to this normalized frequency, with the maximal frequency in black and a null frequency in white. The pink stars represent the values of the dihedral angles for the crystallized Cilengitide (sequence: RGDfV’, PDB ID: 1L5G) in complex with integrin.

For the EGSCyP method, the potential energy of the whole peptide was projected on the diagrams for each residue as a function of the *ϕ* and *ψ* dihedral angles. This energy was computed in kcal/mol with amberff96 force field at the last step of the relaxation with Sander. More precisely, a specific *ϕ*, *ψ* pair (within the grid spacing of 10 degrees), at a given residue position, is shared by a large number of conformations of the whole peptide, having a large distribution in energies. From each of these distributions, we only plotted the minimum energy value. The maps show that the exploration is quite complete, even including conformations of high energy (in blue). The white areas in Figure 3 correspond either to (1) the absence of conformations because of the absence of solution at the IK solver step or of atomic collisions in the proposed structures; (2) conformations of very high energy. Indeed, we considered an energy threshold for a better understanding of the diagrams at −235kcal/mol.

##### Coverage percentage

We defined the coverage of the diagrams as the percentage of *ϕ*-*ψ* pairs, within the grid spacing of 10°, for which at least one conformation was found by EGSCyP. Here the ARG and VAL residues have been chosen as exhaustively sampled residues, with 97% and 98% of coverage level, while the other three are sampled by IK, with coverage levels of 86% to 93%, confirming the fact that exhaustive sampling produces a larger coverage of the diagrams.

One may note that the coverage in the diagrams of Figure 3 seems lower due to the energy threshold and to the color gradient from blue to white. Details about the coverage percentage are available as supplementary material in Table S5. This result shows again the completeness of the exploration. For REMD simulations, the normalized frequency of the observed conformations during the simulation was projected on the *ϕ* and *ψ* axes for each residue. The coverage percentages are this time much lower, varying from 14% of coverage for the D-phenylalanine diagram up to 49% for the arginine diagram. This result is not totally surprising because REMD simulations tend to favor low energy areas and can have some difficulties to cross high energy barriers. In order to evaluate the performance of EGSCyP, we compared the coverage of the Ramachandran plots of Cilengitide for the two methods. For all residues, less than 1% of the areas in the diagrams are visited by REMD, but not by EGSCyP. In other words, 99% of the conformations in the Ramachandran plots produced by REMD are also sampled by EGSCyP. On the contrary, EGSCyP samples more conformations than REMD: the fraction of the map explored only by EGSCyP varies from 48% for glycine up to 79% for valine. This result supports again the good performance of our approach to exhaustively describe the conformational landscape.

##### Map similarities

A visual comparison of the maps clearly indicates that the lowest-energy basins for EGSCyP (in red and black, in the third column of Figure 3) correspond quite well to the high-population areas of the REMD simulations (in red and black too, in the fourth column of Figure 3). Areas with few (or no) conformations found during the REMD simulations, in blue (or white, respectively), correspond essentially to high-energy areas sampled by EGSCyP. Thoroughly analyzing the diagrams, some minor differences between the two methods can be observed. REMD simulations produced only one black region per residue (corresponding to the maxima of frequency) and one or two other regions in red/orange, while EGSCyP resulted in several black points (corresponding to several energy minima). Note however that both maps display different quantities: while the EGSCyP method is based on the potential energy, frequencies derived from the REMD trajectories reflect the total free energy of the system, including the entropic contribution. The difference between the potential energy of the black and red/orange areas is very low with EGSCyP, typically 2kcal/mol. This explains why the valine and phenylalanine diagrams present three black dots in EGSCyP but only one in REMD. These three points in EGSCyP maps correspond to the same energy basin in REMD maps.

#### Comparison of energy minima conformations

##### Clustering

We applied our clustering approach to the results obtained by EGSCyP and REMD simulations for Cilengitide (RGDfV’) in order to compare the energy minima conformations. As a distance metric, we used RMSD based on the *ϕ*-*ψ* dihedral angles. In general, this provides better results than C_*α*_-RMSD for clustering conformations of peptides. In particular, angular RMSD is very useful to identify peptide-plane flips^73^: a large amplitude rotation around the peptide plane affecting the values of the *ψ*_*i*_ and the *ϕ*_*i*+1_ dihedral angles. Indeed, the variations of these two angles mutually compensate, such that the RMSD computed on the *α* carbons does not show any fluctuation^74^. However, such a perturbation can affect the conformation of the peptide, in particular by restraining the side chains conformation. The distance threshold for clustering was set to 60°, which allows distinguishing important peptide-plane flips.

We obtained 120 clusters with EGSCyP against 12 clusters with REMD simulations for Cilengitide. This huge difference is once again not surprising and agrees with the previous results (percentage coverage of Ramachandran diagrams), demonstrating the exhaustiveness of the EGSCyP exploration. The diversity of the clusters are presented as supplementary materials (Figures S6 and S7).

With REMD, the first and most populated cluster represents 57% of the conformations found during the simulation. The second cluster represents 42%. This means that 99% of the conformations are described by these two clusters. We extracted from the trajectory the two structures representing the centers of these clusters and minimized them with Sander. We will call the minimized conformations 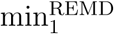 and 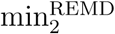 in the following. Their energies are very similar: −267.6 kcal/mol and −267.9 kcal/mol, respectively. Representative conformations of the two first clusters obtained with EGSCyP were also locally minimized without any restraint. The energies of these two minima, called 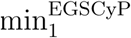 and 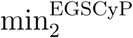, are −267.3 kcal/mol and −267.4 kcal/mol, respectively. As for the two main minima obtained from REMD, the energy values of the minima for EGSCyP are extremely similar. This is in agreement with the fact that peptides, unlike the proteins, present several energetically equivalent minima. Note that the energy values for the minima obtained by the two methods are also very close. Figure 4 shows the superimposed structures of these four minima: in Figure 4-(a), 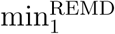 (in green) and 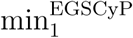 (in beige); in Figure 4-(b), 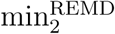 (in pink) and 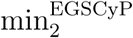 (in blue). The backbones show an important similarity between both methods: 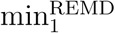 and 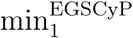 have the same distribution of dihedral angles with an angular RMSD of 32° between them. The second clusters, 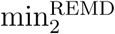 and 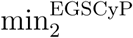, also correspond to each other with a RMSD of 32°. The values of the dihedral angles are presented as supplementary materials (Table S6). The most significant difference between the two groups (Figures 4-(a) and 4-(b)) corresponds to a peptide-plane flip between *ψ*_*I*_ and *ϕ*_*II*_ angles. This difference between the two main minima is illustrated in Figure 4-(c). The structures of 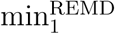 and 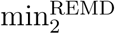 are represented with carbon atoms in green and pink respectively. The flip explaining the difference is surrounded in red. It impacts the position of the arginine side chain. Indeed, in 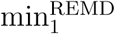, the side chain can fold and create a hydrogen bond with the side chain (in green) of the aspartate. Considering 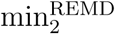, the side chain (in pink) is oriented more outwards and makes a hydrogen bond with the oxygen of the valine carbonyl group. The comparison of the two conformations shows that the flip leads to a tilt of the carbonyl group of the glycine visible on the onset of Figure 4-(c). This prevents the arginine side chain of the second minimum to come closer to the cycle and interact with the aspartate side chain. Indeed, the orientation of the oxygen atom would create a collision if the side chain would attempt to get closer (see van Der Waals radius of the carbonyl oxygen atom in the zoom of Figure 4-(c)). Therefore, the tilt has an impact on the structure, constraining the arginine side chain (even if it does not mean that all the structures from the cluster corresponding to 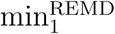 have this side chain orientation).

**Figure 4:**
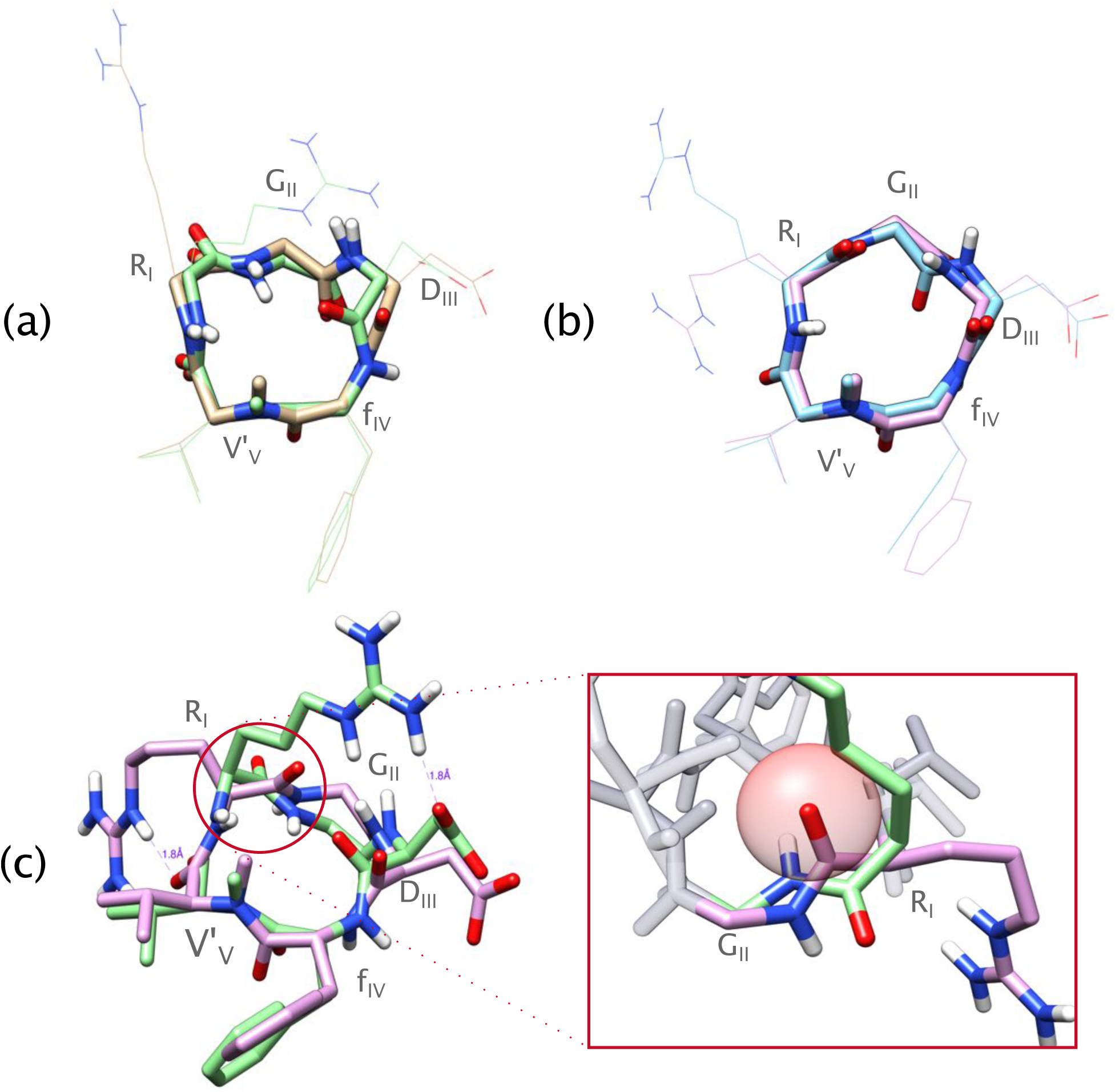
Lowest energy conformations obtained for Cilengitide (RGDfV’) with EGSCyP and REMD. (a) Superimposition of the structures of 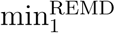 (in green) and 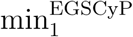 (in beige). RMSD on dihedral angles between both structures equals 32°. (b) Superimposition of the structures of 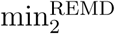 (in pink) and 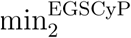 (in blue). RMSD between the two structure equals 32^°^. For (a) and (b) the side chains are represented in lines for better visibility. (c) Superimposition of the structures of 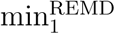 (in green) and 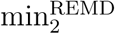 (in pink). The purple dotted lines represent the measurement of the distance between two atoms. The major difference between the two clusters is a peptide flip between the arginine and glycine surround in red. On the right part of the figure, a zoom was made on the flip from the other side for a better visibility. The red sphere represents the van Der Waals radius of the carbonyl oxygen atom from 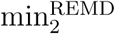.

##### Comparison with crystallographic structure

Results provided by the two conformational exploration methods were compared to the X-Ray structure of Cilengitide bound to integrin (PDB code: 1L5G), which, to the best of our knowledge, is the only available structure of this peptide. The dihedral angles values of this X-Ray structure were projected on all the RGDfV’ Ramachandran diagrams. They are indicated with a pink star in the third and forth columns of Figure 3. We can observe that this experimental structure is close to energy minima (in black). The RMSD on dihedral angles was computed between this structure and the clustered minima structures from the two methods (see section above). The 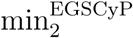 minimum is the closest one from the X-Ray structure, with RMSD equal to 14°. The RMSD for 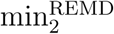 is 35°. The first minima (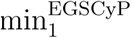 and 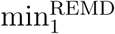) are more distant to the experimental structure, with RMSD around 75°. In Figure 5, the experimental structure (in purple) is superimposed to 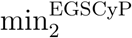 (in blue) in order to show the good similarity of the two conformations. However, it can be noticed that the experimental peptide is bound to the protein and not free.

**Figure 5:**
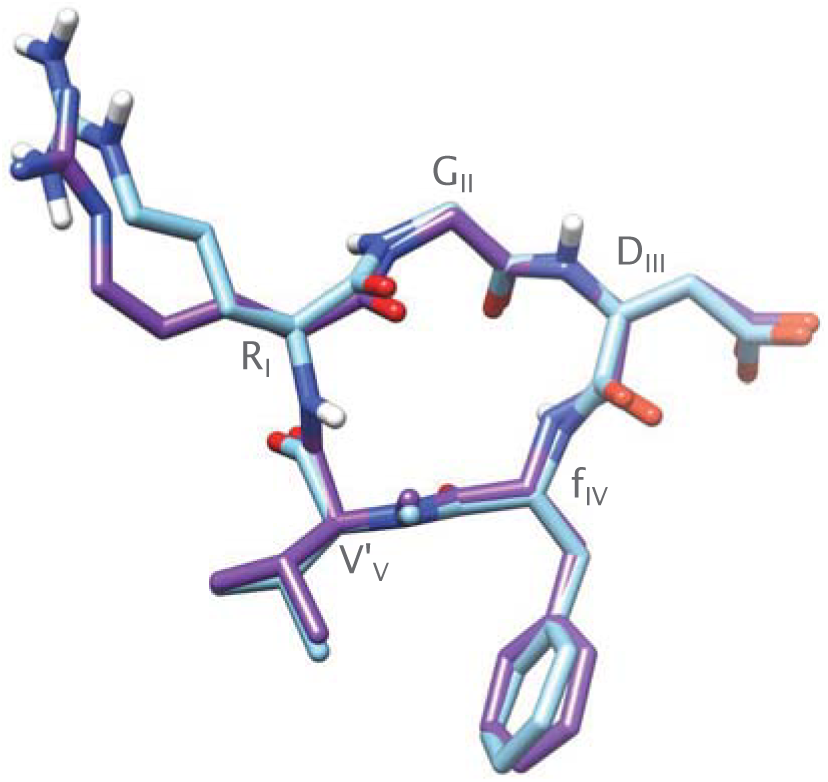
Structure of Cilengitide (RGDfV’). Comparison between the crystallographic structure (PDB ID: 1L5G, in purple) and the closest energy minimum found by EGSCyP (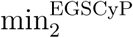 in blue). RMSD of the dihedral angles between them is 14°.

#### Side chain placement

Figure 6 represents the projections of the energy landscape of the RGDFV peptide as a function of the *χ*_1_ and *χ*_2_ dihedral angles^75^ for the arginine, aspartate and phenylalanine residues. The first column corresponds to the conformations obtained with the EGSCyP using SCRWL4 for the side chain positioning. The second column corresponds to the EGSCyP with the alternation of BH and local minimization, referred to as BH/MC in the following. The last column corresponds to the conformations from the REMD simulations. The color code corresponds to the potential energy of the whole peptide for the EGSCyP and to the normalized frequency of the conformations found during the simulations at the lowest temperature replica for the REMD simulations.

**Figure 6:**
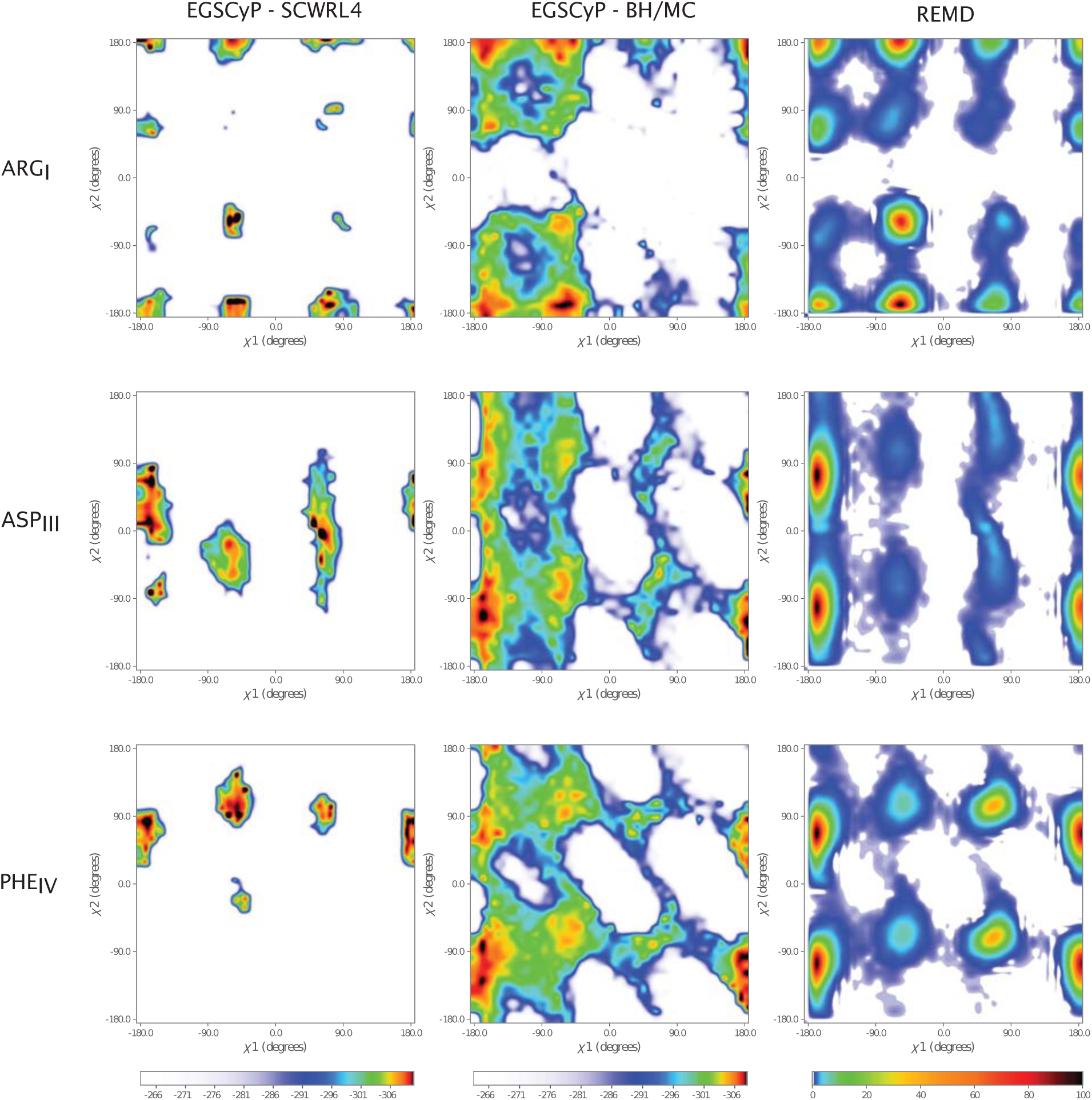
Comparison of the side chain conformational landscapes obtained for the RGDFV cyclic peptide with EGSCyP using SCWRL4^49^ (left column), with EGSCyP and the alternation of BH and local minimization (middle column), and with the REMD simulations (right column). The conformational landscape was projected as a function of the *χ*_1_ *χ*_2_ dihedral angles of the side chains for ARG (first line), ASP (second line) and PHE (last line). The diagrams were created using the matplotlib python library^78^. For EGSCyP, the color codes correspond to the minimal potential energy of the whole peptide associated to this combination of *χ*_1_-*χ*_2_. For the REMD simulations, the color code corresponds to the normalized frequency 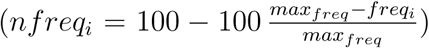 of the conformations found during the simulations at the lowest temperature replica as a function of *χ*_1_ and *χ*_2_.

The *χ*_1_ - *χ*_2_ maps show that there is a striking difference between the two methods for the side chain placement. With SCWRL4, the covered space is very small compared to the one obtained with BH/MC or the REMD simulation. Actually, SCWRL4 covered 8% to 16% of the diagrams while the BH/MC covered between 54% and 69%. Less than 1% of the space is covered only by SCWRL4 and not by BH/MC meaning that the space explored by SCWRL4 is almost totally included in the one explored by BH/MC. Regarding the quality of the obtained conformations, the energy minima obtained with SCRWL4 are not localized at the same place as the frequency maxima obtained by REMD. For the arginine residue, the minima are rather close to the frequency maxima for the arginine obtained by REMD, but for the aspartate and phenylalanine, they are not localized on the areas of high frequency of the REMD simulations. On the contrary, the BH/MC sampling shows a good correspondence between its energy minima basins and the frequency maxima from the REMD simulations for the three residues of Figure 6. The percentage coverage for the REMD side chain placement is rather constant among the different residues (around 58%) and the part of the coverage in common between the BH/MC and REMD is about 50% of the diagrams. Actually, less than 10% of the coverage in the REMD diagrams are not covered by BH/MC. Therefore, this study confirms that the use of SCWRL4 is definitely not adapted for the case of small cyclic peptides. Moreover, the method cannot deal with chemical modifications. These results also validate the good performance of the alternation BH/MC for side chain placement within EGSCyP. The *χ*_1_ - *χ*_2_ diagrams for the other three peptides are available in supplementary materials (Figure S8). Similarly, they show a good agreement between BH/MC and REMD coverage of the side chain maps (Table S7), in particular with the experimental structure of Cilengitide.

### Effects of the chemical modification on the energy landscape

The exploration of the conformational landscape of the three RGD peptides (RGDFV, RGDfV and RGDfV’), was compared to analyze the influence of the chemical modifications, as it can be observed in Figure 3.

#### D-residues

The conformational landscape of the D-residue (noted as a lower case letter) changes compared to its L-enantiomer, as expected. The accepted conformations show some symmetry relative to the centers of the Ramachandran plots of Figure 3. There are three energy minima basins for the L-enantiomer, but only one for the D-enantiomer (black dots in the first and second columns of Figure 3). Nevertheless, the major basin for phenylalanine (upper left for the L-enantiomer) is conserved through symmetry in the D-enantiomer. Therefore, the landscape of the D-residue seems more restrained than the one of the L-enantiomer, confirming the properties of the D-residues to locally constrain the conformation of the molecule^2^. Actually, the percentage of coverage is quite similar between the two diagrams (about 93%), but the potential energy is globally higher with the D-residue (blue areas).

The replacement of the L by the D-enantiomer of the phenylalanine has an impact on the landscapes of the whole peptide, extending to the second neighbors. This can be observed both on the level of the covered surface and the number of basins. The smaller number of basins within the peptide containing one L amino acid suggests a higher global constrained conformation when a non natural amino acid is included. The two direct neighbors are particularly impacted: for the downstream valine, one (around *ϕ* = 70 and *ψ* = −70) of the two energy minima basins vanishes and the area *ϕ* > 0 has conformations of higher energies. For the upstream aspartate, the basin goes from the lower part of the diagram (*ψ* < 0) to the upper part (*ψ* > 0) and the area with *ψ* < 0 is of higher energies. For the second neighbor, differences can still be shown: the glycine energy basin from the lower left part of the diagram vanishes, the one from the lower right part is much less extensive, and the upper part of the diagram (*ψ* > 0) presents higher energies when the L-phenylalanine is used. For the aspartate, the differences are the following: the energy basin from the lower part of the diagram disappears and the one from the upper part shrinks; the right part of the diagram also presents higher energies with the D-enantiomer. Therefore, these remarks confirm that the D-enantiomer does not only constraint the conformational landscape of the peptide, but also affects its direct and indirect neighbors.

#### N-methylation

The consequence of the addition of a N-methylation on valine can be evaluated by comparing the RGDfV and RGDfV’ peptides (Figure 3, column 2 and 3). The coverage percentage of the landscape is quite constant, even in spite of the increasing number of collisions. Inspecting the diagram for each residue, one can make the following observations. 1) The presence of the methyl on the backbone shifts the energy towards higher values at the location of the modification, so it is consistent with some kind of destabilizing effect of this modification. 2) The central zone of the Ramachandran plot, highly unfavorable, is enlarged for the modified residue; this does not significantly affect the other residues. 3) The number of basins goes from one for the natural valine up to three for the modified one. 4) The diffusing effect of the methylation occurs on the landscape of arginine, the first neighbour in the cycle. One recovers the basins in the lower left of the ARG diagram that disappeared under the presence of the D-phenylalanine. Therefore, there is a sort of balance between the constraints introduced by the D-form and the methylation that increase the number of basins. Thus, one may hypothesize that the addition of the N-methyl offsets a part of the constraints brought by the D-enantiomer.

## Conclusions

Small cyclic peptides present unique properties making them promising therapeutics drugs. The current difficulty to predict the structure and to design cyclic peptides could be greatly improved thanks to a better understanding of their whole conformational landscape. In this paper, we propose a method, called EGSCyP, for the exhaustive exploration of the energy landscape of cyclic pentapeptides possibly involving chemical modifications. We have shown the good performance of the method, which is based on a robotics approach and a multi-level representation of the peptide. The comparison of the results obtained for three cyclic pentapeptides with REMD simulations reveals the completeness and consistence of our approach. Also, the comparison with the experimentally-determined structure of Cilengitide bound to the integrin complex demonstrates the predictive capabilities of the method. Moreover, we have demonstrated the effectiveness of the alternation of BH/local minimization for the sampling of the side chains conformations, whereas SCWRL4 fails to correctly position the side chains of small cyclic peptides. Finally, this approach clearly shows the effect on the conformational landscape of the D-residue that constraints the landscape and the N-methylation that also modulates it.

In the short future, we will apply EGSCyP to a larger dataset. We will also investigate conformational changes using the connectivity graph built during the exploration. Another interesting perspective would be to use force fields optimized for cyclic peptides. For instance, the RSFF1 and RSFF2 force fields have shown good performance on cyclic peptides^76^. However, progress is still needed, since it has been shown that these force fields are for the moment not well suited to the use of non natural residues like the N-methylated ones^77^ neither for implicit solvent. The better understanding of the conformational properties of small cyclic peptides may be used to develop more suitable methods for structure prediction and design. Finally mention that we intend to extend the approach to larger cyclic peptides. Obviously, a systematic, grid-based exploration as described in this paper has limits in the length of the peptide candidates due to the combinatorial explosion. However, the proposed multi-level modeling approach can be exploited within stochastic exploration-optimization methods, such as variants of BH, able to provide a global picture of the conformational landscape.

## Acknowledgments

This work was performed using HPC resources from GENCI-CINES (Grant 2016-c2016077641). CALMIP is gratefully acknowledged for the access to the super-computer EOS for developing the EGSCyP method. We also thank Dr Michele Lazzeri for letting us using his cluster, and Dr Guillaume Postic for valuable discussions and advices.

## Abbreviations

BH: : Basin Hoppin
DoF: : degree of freedom
EGSCyP: : Exhaustive Grid Search for Cyclic Peptides
IK: : Inverse Kinematics
PDB: : Protein Data Bank
MC: : Monte Carlo
REMD: : Replica Exchange Molecular Dynamics
RMSD: : Root Mean Square Deviation

